# One-time Nitrogen Fertilization Shifts Switchgrass Soil Microbiomes within a Context of Larger Spatial and Temporal Variation

**DOI:** 10.1101/520239

**Authors:** Huaihai Chen, Zamin K. Yang, Dan Yip, Reese H. Morris, Steven J. Lebreux, Melissa A. Cregger, Dawn M. Klingeman, Dafeng Hui, Robert L. Hettich, Steven W. Wilhelm, Gangsheng Wang, Frank E. Löffler, Christopher W. Schadt

**Affiliations:** Biosciences Division, Oak Ridge National Laboratory; Department of Biological Sciences, Tennessee State University; Chemical Sciences Division, Oak Ridge National Laboratory; Environmental Science Division, Oak Ridge National Laboratory; Department of Microbiology, University of Tennessee; Institute for Environmental Genomics and Department of Microbiology & Plant Biology, University of Oklahoma

## Abstract

Soil microbiome responses to short-term nitrogen (N) inputs within the context of existing spatio-temporal variability remain uncertain. Here, we examined soil bacterial and fungal communities pre/post-N fertilization in an 8 year-old switchgrass field, in which twenty-four plots received N fertilization at three levels (0, 100, and 200 kg N ha^-1^ as NH_4_NO_3_) for the first time since planting. Soils were collected at two depths, 0-5 and 5-15 cm, for DNA extraction and amplicon sequencing of 16S rRNA genes and ITS regions, and soil metagenomic analysis. Baseline assessment prior to fertilization revealed no pre-existing differences in either bacterial or fungal communities across plots. The one-time N fertilization increased switchgrass yields and tissue N content, and the added N was nearly completely removed from the soil of fertilized plots by the end of the growing season. Both bacterial/archaeal and fungal communities showed large spatial (by depth) and temporal variation (by season) within each plot, accounting for 17 and 12-22 % of the variation in bacterial/archaeal and fungal community composition, respectively. While N fertilization effects accounted for only ~4% of overall variation, some specific microbial groups, including the bacterial genus *Pseudonocardia* and the fungal genus *Archaeorhizomyces,* were notably repressed by fertilization at 200 kg N ha^-1^. Bacterial groups varied with both depth in the soil profile and time of sampling, while temporal variability shaped the fungal community more significantly than vertical heterogeneity in the soil. Thus, variability within the field might override the changes induced by N addition. Continued analyses of these trends over time with fertilization and management are needed to understand whether these transient effects change over time.

## Introduction

Cultivation of dedicated bioenergy crops is of interest to sustain long-term energy supplies [1]. The International Energy Agency predicts that biofuels could satisfy more than a quarter of world needs for transportation energy by 2050 [2]. Switchgrass *(Panicum virgatum* L.) has been a prominent candidate as an energy crop due to its high biomass yield, low maintenance and limited-input requirements [3], and high adaptability to marginal sites [4]. Such characteristics may allow switchgrass for its use to reclaim degraded or abandoned agricultural lands while reserving fertile lands for food production [5]. With its well developed and deep rooting systems, switchgrass may also improve belowground carbon storage and nutrient acquisition [6] and potentially moderate the diversity of below-ground and plant-associated microbiomes. Thus, how switchgrass cultivation affects soil microbial communities and their interaction with crop yields needs further investigation to understand the long-term ecosystem consequences and sustainability of the cultivation of perennial crops, such as switchgrass.

Soil microbial communities play fundamental roles in terrestrial ecosystems, such as regulating the decomposition of organic matter as well as driving nutrient cycles and energy flow [7, 8]. To this end, these microbiomes have considerable effects on soil quality and agricultural sustainability [9]. However, soil management with fertilizer additions may shift soil microbial abundance and composition as well as functions by affecting soil physical and chemical characteristics [10]. For example, laboratory studies showed that N addition depresses soil microbial activity, microbial biomass, and enzyme activities by shifting the metabolic capabilities of soil bacterial communities toward the decomposition of more labile soil carbon pools [11]. In addition, nutrient inputs have been shown to shift the composition of soil microbial communities in consistent ways in grasslands across the globe with reduced average genome sizes of microbial communities following nutrient amendment, leading to decreased relative abundances of some important microbial functional groups, such as methanogenic archaea, oligotrophic bacteria and mycorrhizal fungi [12]. Several reasons may account for such microbial responses to fertilization. Fertilization may cause soil acidification, and thus alter soil microbial diversity and composition [10, 13]. Additionally, nutrient amendments may have direct effects on organic matter decomposition, leading to changes in the quantity and quality of resources available for microbes, and therefore reshape microbial community structure based on their substrate utilization preferences [14]. However, our understanding of the mechanisms of N fertilization effects on microbial communities are mostly based on long-term fertilization, in which edaphic soil properties have likely been significantly altered by soil management over time. Although transient nutrient enrichment effects upon terrestrial microbial C and N processes have been reported [15–17], our understanding of the immediate response of the below-ground microbial community to N inputs is still limited, and sometimes inconsistent with results of longterm experimental data [18–20]. Additional research is thus necessary to evaluate microbial community dynamics and their interactions with nutrient cycling under the no-, low-or periodic fertilization regimes that would be optimal for sustainable perennial bioenergy crop production scenarios [21, 22].

Besides soil nutrient availability, soil microbial distribution is influenced by a wide range of soil characteristics, such as soil pH, substrate quantity and quality, moisture and oxygen levels, nearly all of which could typically change with soil depth [23] and vary over seasons [24]. Soil depth and measuring time in a growing season thus influence patterns of spatial and temporal community variation [25, 26]. Compared to top soil, subsurface soils have higher mineral content, less aeration, and lower organic carbon availability. Thus, microbial biomass and diversity typically decrease rapidly with depth in the soil profile [27]. Often, most variability in microbial community composition occurs in surface soils, while deeper soils have more similar communities regardless of soil management [23]. Seasonal variability also has a large influence on microbial communities [28]. For example, seasonal changes of temperature and soil moisture can directly shape microbial communities [29, 30]. Moreover, seasonal changes in plant growth and allocation can indirectly affect soil C inputs [31, 32]. Lauber et al. [24] investigated the temporal variability of bacterial communities in different ecosystems, showing that most of the temporal variation in bacterial composition within an agricultural field could be explained by soil moisture and temperature variations. Given these previous studies, it is possible that the shifting spatial and temporal patterns of soil microbial communities may overwhelm short-term soil nitrogen management effects and needs to be accounted for in such assessments.

Here, we used high-throughput barcoded sequencing to assess short-term effects of one-time N fertilization on the spatio-temporal variation of soil microbial communities in an 8 year-old switchgrass field, over two soil depths and across four sampling seasons. We hypothesized that (1) one-time nutrient inputs could significantly change above-ground plant yields and substrate quality, and re-shape soil bacterial and fungal communities, but that short term N effects would be modest compared to existing spatio-temporal variation, (2) bacterial and fungal composition would differ spatially and temporally, but the response of these communities to the N-fertilization would be taxon specific.

## Materials and Methods

### Site characterization, experimental design and plant and soil sampling

The experiment was established in an eight year-old switchgrass field near the Heritage Center, located on US DOE land, in Oak Ridge, Tennessee, USA (35.9255 N, 84.3947 W). The 10-year mean annual temperature and annual precipitation at the site were 14.1°C and 1436 mm, respectively. The field was in pasture and hay rotations when taken over by DOE in the 1940s during the Manhattan Project. However, due to proximity of floodplain of the Clinch River and Poplar Creek, the land was never developed and instead maintained as wildlife habitat and riparian buffer as a field of mixed grasses and forbs, using a combination of mowing and prescribed burning. In 2009, under contract for UT Institute of Agriculture and Genera Energy, the site was cleared, seeded, and subsequently managed for switchgrass production. After the contract expired in 2012, the site remained in switchgrass, but has again been managed as buffer and wildlife habitat, and maintained only with periodic prescribed fire and mowing. In mid-December 2016, the switchgrass field was mowed to a 10-cm stubble height and twenty four plots (5 m × 5 m) were set up including three N fertilization levels (0, 100, and 200 kg N ha^-1^) with eight replicates based on a complete randomized design (Fig. 1). A 2.5-m inter-plot “alley way” was periodically mowed to allow access and separate the plots between treatments and replicates. Just before spring emergence of the switchgrass (March 30, 2017) commercial ammonium nitrate (34% nitrogen) was hand-applied to fertilizer-treated plots with N fertilization levels of 100, and 200 kg N ha^-1^. Post-emergence (June 20, 2017), all plots were treated with Garlon 3A herbicide as prescribed by the manufacture to help control broadleaf weeds. After fall senescence (November 13, 2017) above-ground biomass of switchgrass was measured [33] using a sickledrat to harvest all aboveground biomass from a 0.1 m^2^ area at four randomly chosen locations in each plot. The four samples of aboveground biomass were pooled by plot in paper bags, oven dried at 70^°^ C, and weighed to determine dry mass per unit area. A subsample of the plant material was then ground into powder using a laboratory mill before total C and N were determined by dry combustion method using a Perkin-Elmer 2400 CHN analyzer (Perkin-Elmer Corporation, Norwalk, CT, USA).

**Fig. 1.**
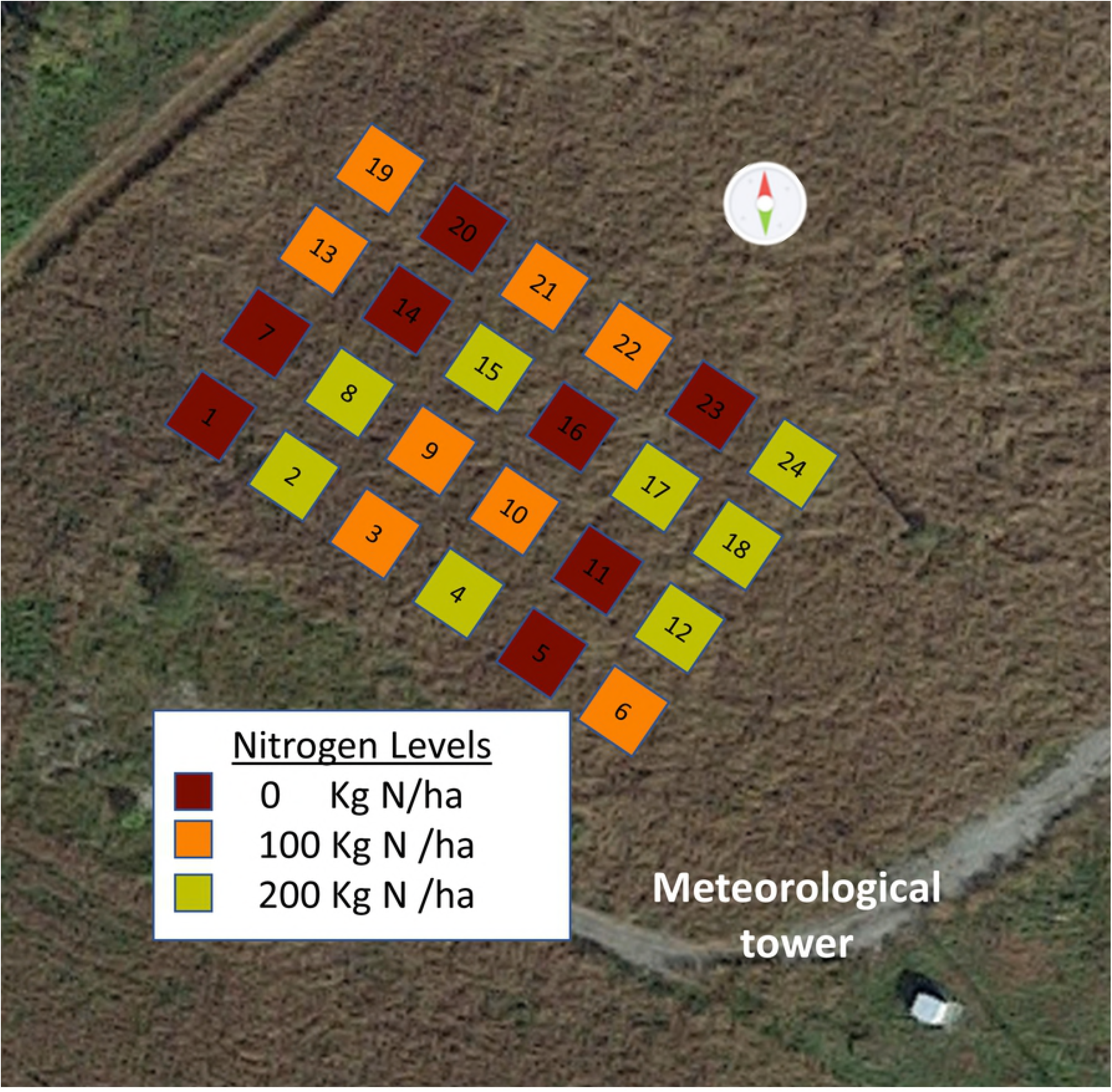
A Google map showing twenty four plots (5 m × 5 m) of three N fertilization levels (0, 100, and 200 kg N ha^-1^) with eight replicates based on a complete randomized design.

For soil DNA and chemical characterization, four sets of soil samples were collected across seasons including Winter 2016 (December 16, 2016), Spring 2017 (April 5, 2017), Summer 2017 (July 5, 2017), and Late Fall 2017 (November 15, 2017). At each sampling event, soil cores (2.5 cm diameter ×15 cm height) were collected randomly from each plot and separated into two depth increments of 0-5 and 5-15 cm. Soils collected in Winter 2016 (before N fertilization) were used to assess soil microbiomes and metagenomes, and check whether there was systematic preexisting differences of microbial communities across plots. Soils collected following N fertilization, *i.e.,* Spring 2017, Summer 2017, and Late fall 2017, were used to compare difference in microbial communities among N treatments. Soil collected for all four seasonal samplings was used to assess how soil depth and sampling seasons affected microbial communities across all three nitrogen treatments. All soil samples for microbial analyses were transported on dry ice to the lab and stored at -80^°^ C prior to soil DNA extraction. Total C and N were determined on samples collected in the Summer of 2017 using the dry combustion method and soil inorganic N (NH_4_^+^-N and NO_3_^-^-N) was analyzed using a FIA QuikChem 8000 autoanalyzer (Lachat Instruments, Loveland, CO, USA). Generally, 0-5 cm soils had 4.5% soil total C, 0.2% soil total N, 10.2 mg kg^-1^ NH_4_^+^, and 1.3 mg kg^-1^ NO_3_^-^, while 5-15 cm soil had significantly lower 1.8% soil total C, 0.03% soil total N, 1.8 mg kg^-1^ NH_4_^+^, and 0.3 mg kg^-1^ NO_3_^-^.

### DNA extraction, rRNA gene amplicon sequencing, and metagenomic sequencing

Approximately 10 g of soil from each sample was homogenized in a mortar and pestle with liquid N_2_, and soil DNA was extracted from a 0.25 g aliquot of the soil sample using the MoBio DNeasy PowerSoil Kit (Qiagen, Carlsbad, CA, USA) according to the manufacturer’s instructions. DNA concentrations were determined and purity was confirmed by the ratio of absorbance at 260 and 280 nm (1.70-1.90) using a NanoDrop 1000 spectrophotometer (NanoDrop Products, Wilmington, DE, USA).

A two-step PCR approach was used to barcode tag templates with frameshifting nucleotide primers for amplicon sequencing [34] with some modifications previously described [35]. To increase phylogenetic coverage for community analysis of bacteria, archaea, and fungi, a group of nine forward and six reverse primers for bacteria and archaea, and another group of eleven forward and seven reverse primers for fungi, mixed at equal concentration of 0.5 μM were used to target 16S rRNA V4 region and fungal ITS2 rRNA, respectively [35]. Primary PCR was conducted for 5 cycles of 1 min at 95 ^°^C, 2 min at 50 ^°^C, and 1 min at 72 ^°^C, followed by a final elongation of 5 min at 72 ^°^C. This PCR product was then cleaned up using Agencourt AMPure beads (Agencourt Bioscience, Beverly, MA, USA) and eluted in 21 μL of nuclease-free water.

To tag amplicons with barcoded reverse primers and forward primers, 20 μL of purified DNA fragments from the primary PCRs were added to 50 μL secondary PCR assays, which were initiated at 95 ^°^C for 45 sec, followed by 32 cycles of 15 sec at 95 ^°^C, 30 sec at 60 ^°^C, and 30 sec at 72 ^°^C, followed by a final elongation of 30 sec at 72 ^°^C. The use of separate tagging reactions can help reduce heterodimers because PCR clean-up more efficiently removes shorter primers [34]. Up to ninety six secondary PCR products were then pooled based on agarose gel band intensity, followed by a second clean-up with Agencourt AMPure beads (Agencourt Bioscience, Beverly, MA, USA) using 0.7-1 of bead-to-DNA ratios. The mixtures of the purified 16S rRNA gene or ITS amplicon fragments were then paired-end sequenced on Illumina Miseq platform (250×2 paired end, v2 chemistry) (Illumina, San Diego, CA, USA) using a 9 pM amplicon concentration.

To examine microbial community potential and function prior at the site, DNA extracted from soil samples collected in Winter 2016 were pooled to form four composite samples. Specifically, for each of two soil depths, DNA from No. 1-12 and 13-24 plots was pooled together, respectively (two depth × two replicates). Shotgun metagenomes were prepared using Nextera XT sequencing libraries (Illumina, San Diego, CA) according to the manufacture’s recommendations using 500ng of DNA (15031942 v03). Final libraries were validated on an Agilent Bioanalyzer (Agilent, Santa Clara, CA) using a DNA7500 chip and concentration was determined on an Invitrogen Qubit (Waltham, MA) with the broad range double stranded DNA assay. Barcoded libraries were pooled and prepared for sequencing following the manufactures recommended protocol (15039740v09, Standard Normalization). One paired end sequencing run (2 x 300) was competed on an Illumina MiSeq instrument (Illumina, San Diego, CA) using v3 chemistry.

### Bioinformatic and Statistical Analyses

Forward and reverse primers were trimmed with Cutadapt [36]. Paired-end sequencing data were then joined and demultiplexed using QIIME [37] with quality filter at Phred > 19. Chimeras of trimmed and filtered sequences were identified and removed using a usearch method in QIIME. Operational taxonomic units (OTUs) with 97% identity were picked with the open reference algorithm and usearch61 otu-picking method. Taxonomy was assigned using the RDP (Ribosomal Database Project) taxonomy-assignment method [38] against the most recent version of Greengenes database (13.8) for 16S rRNA sequencing data and UNITE database (12.11) for ITS sequencing data. All global singletons were removed from the dataset. The 16S and ITS OTUs were further analyzed for alpha and beta diversity using QIIME. Metrics for analyzing beta diversity were Bray-Curtis distance. Bacterial community functional traits were predicted using PICRUSt (Phylogenetic Investigation of Communities by Reconstruction of Unobserved States) [39]. The Miseq sequences were deposited on NCBI Sequence Read Archive (SRA) database under the BioProject accession number of PRJNA512218.

Soil metagenome sequences were uploaded to Rapid Annotation using Subsystems Technology for Metagenomes (MG-RAST; http://metagenomics.anl.gov) [40] under project accession number mgp22000, and annotated using the RefSeq database for taxonomic assignment and the SEED Subsystems database for functional classification (maximum e-value cutoff was 1e^-5^, minimum identity cutoff was 60%, and minimum alignment length was 50).

One-way analysis of variance (ANOVA) of a completely randomized design (SAS 9.3, SAS Institute Inc. Cary, NC, USA) was used to assess significant differences in above-ground yields and plant C/N contents among N fertilization levels. A three-factor ANOVA of a completely randomized design was used to analyze microbial alpha diversity and the abundances of microbial taxonomic groups among the three N fertilization levels, two soil depths, and four seasonal samplings. Microbial beta diversity was compared using a three-factor PEMANOVA method (N fertilization levels, soil depths, and sampling season) with 9999 permutations conducted in PRIMER (Plymouth Routines in Multivariate Ecological Research Statistical Software, v7.0.13, PRIMER-E Ltd, UK). A RELATE analysis was also performed to evaluate the relatedness between bacterial and fungal beta diversity by calculating Spearman’s Rho correlation coefficient in PRIMER. The DistLM (distance-based linear model) function in PRIMER was used to evaluate the associations of above-ground yields and plant C/N contents with bacterial and fungal beta diversity [41]. Heat maps were constructed using HeatMapper [42] to represent all taxonomic groups at genus level that differed significantly (P < 0.05) among three N fertilization levels, two soil depths, and four sampling times. Venn’s diagrams were also constructed to visualize how many significantly affected bacterial/archaeal and fungal genera were shared between the factors of soil depth and sampling time using Venny 2.1.0 [43]. Additionally, Pearson’s correlation coefficients were examined to further evaluate relationships between the relative abundances of taxa and N fertilization rates.

## Results

### Spatial variation in microbial community structure and function pre-nitrogen addition

Both 16S rRNA gene and ITS region amplicon sequencing revealed no significant pre-existing differences in alpha or beta diversity in either the bacterial/archaeal or fungal communities across the 24 plots in this switchgrass field before N fertilization, however diverse bacterial/archaeal and fungal taxa were observed (Fig. 2). Bacterial communities varied significantly by depth (P < 0.05) with the 0-5 cm soil layer having greater Planctomycetes (8%), Bacteroidetes (7%), and Verrucomicrobia (5%), but less abundant Proteobacteria (34%), Chloroflexi (5%), and Gemmatimonadetes (2%) than the deeper layers (Fig. 2). Surprisingly, fungal phyla did not show any differences between the soil depths examined in these switchgrass soils.

**Fig. 2.**
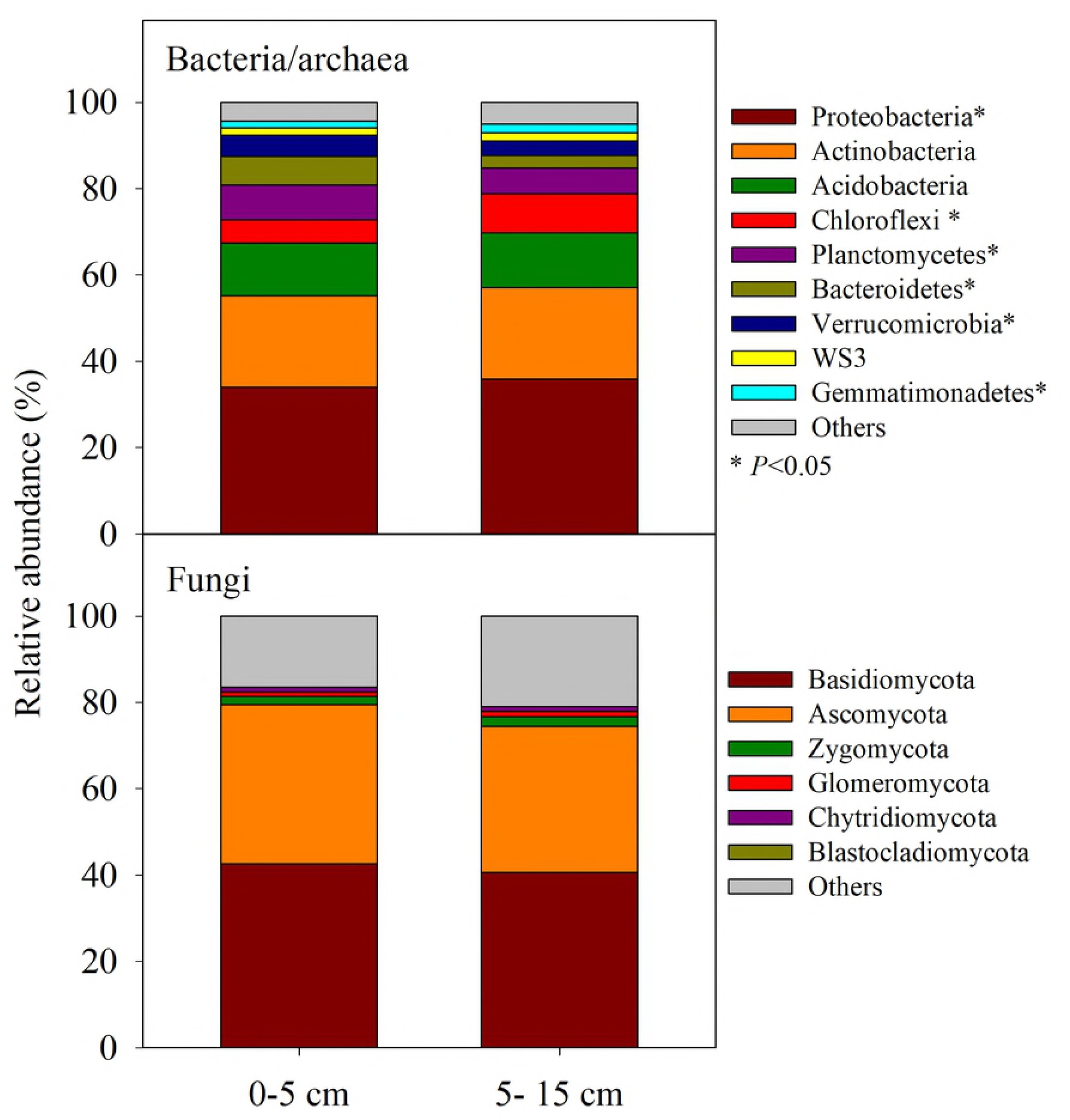
Relative abundances of bacterial/archaeal and fungal dominant phyla (average abundances > 1%) affected by two soil depths (0-5 and 5-15 cm). Asterisks indicate significant difference at α = 0.05 between two soil depths.

Shotgun metagenomes also showed high taxonomic and functional diversity in the switchgrass field (Fig. S1). However, when phylogenetic assignments of the metagenome reads were compared to the relative abundance in 16S rRNA gene amplicon analyses, soil metagenomes indicated significant differences in the datasets accross several of the dominant phyla. There was a 45% increase in Proteobacteria, a 15-fold increase in Firmicutes, and a 2-fold increase in Cyanobacteria in the shotgun metagenomes when compared to 16S rRNA gene amplicon analyses. Other phyla, such as Acidobacteria, Planctomycetes, and Chloroflexi, were reduced by 52-63% in soil metagenomes when compared to 16S rRNA gene amplicon analyses from the same samples and dates. Soil metagenome predicted functional gene profiles were compared to those predicted from PICRUST-based analysis of 16S rRNA gene amplicon data and indicated significantly different profiles (Fig. S2). As a result, PICRUST-based analyses of seasonal functional gene patterns and responses to fertilization were not pursued further.

### Microbial alpha and beta diversity post-nitrogen addition

Although neither bacterial/archaeal nor the fungal alpha diversity were significantly affected by N fertilization levels, both the community richness (Chao1 index) and diversity (Shannon index) showed significant spatio-temporal changes (P < 0.05) (Fig. 3). Between the two soil depths, the 0-5 cm layer had significantly higher Chao1 richness and Shannon evenness indices in both the bacterial/archaeal and fungal communities compared to the 5-15 cm layer (P < 0.05). In analyses of seasonal variation, Chao1 diversity showed a similar pattern. Spring 2017 had lower richness in the bacterial/archaeal community, while Winter 2016 and Fall 2017 had significantly greater richness in fungal communities (P < 0.05). Shannon diversity indices showed significant divergence across seasons (P < 0.05), and the bacterial/archaeal community was more evenly distributed in Fall 2017, whereas the fungal community was more uneven in Summer and Fall 2017 than the other two sampling seasons (P < 0.05).

**Fig. 3.**
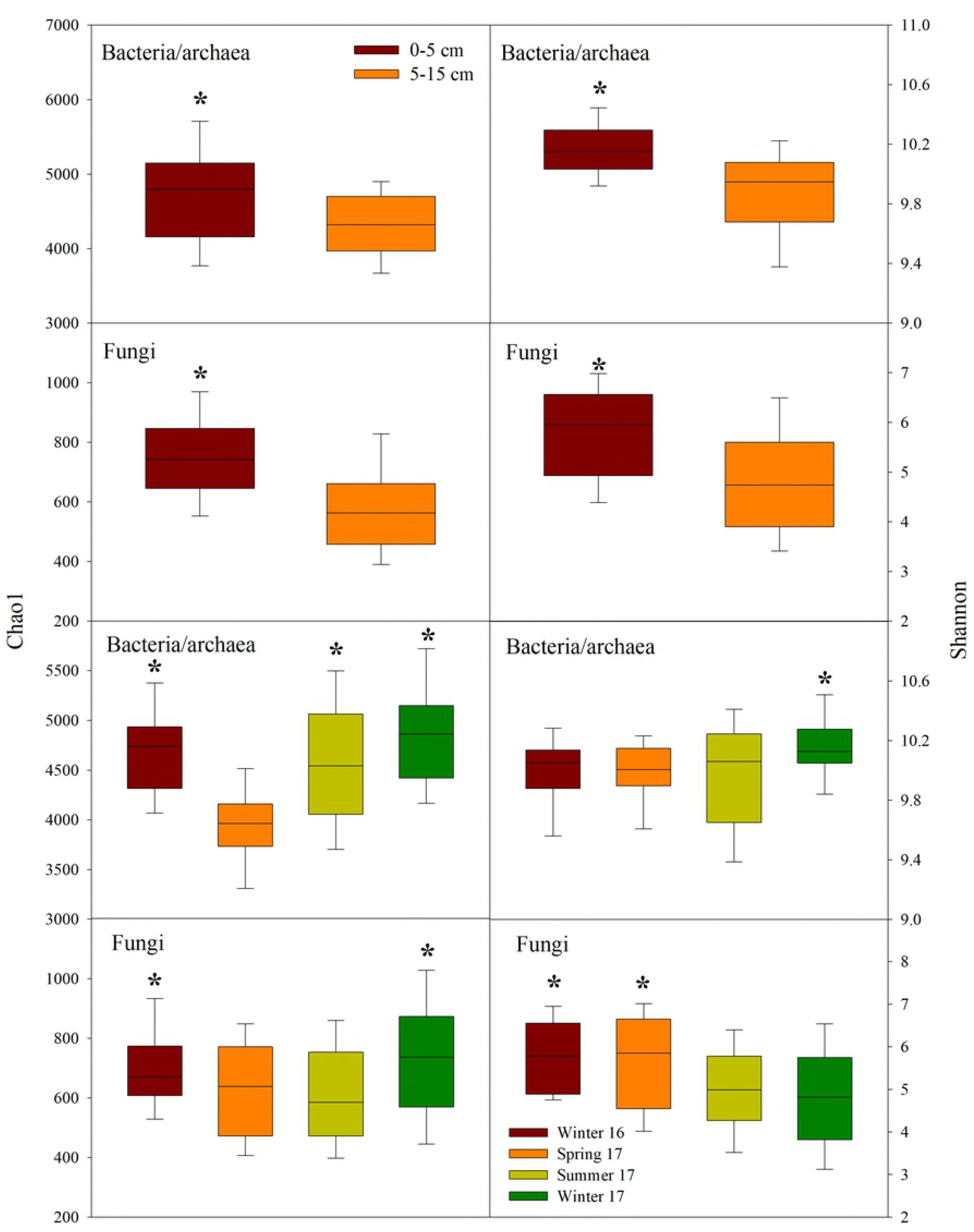
Box plots showing Chao1 richness and Shannon diversity of bacterial/archaeal and fungal communities affected by two soil depths (0-5 and 5-15 cm) and four sampling seasons (Winter 2016, Spring, Summer, and Fall 2017). Sequence depths were 10000 for 16S and 5000 for ITS. Asterisks indicate significant difference at α = 0.05 between two soil depths or among four sampling seasons.

Permanova tests showed that short-term application of N fertilizers caused significant variation in bacterial/archaeal and fungal community composition (P < 0.05) (Table 1 and Fig. 4). Together, N fertilization effects could explain 3.4% of variation in bacterial/archaeal and 4.4 % of fungal community variation (Table 1). However, the spatio-temporal variation (depth and season) were more significant than N effects for bacterial, archaeal and fungal communities (P < 0.0001) (Table 1 and Fig. 4). Soil depth and sampling season contributed to approximately 16.8 and 17.3% of bacterial/archaeal community variation, respectively, and 12.4 and 22.4 % of fungal community change, respectively (Table 1), thus indicating relatively slight short-term effects of N fertilization on microbial communities when compared to the spatio-temporal variation. In addition, RELATE analyses further confirmed that bacterial/archaeal community structures were significantly related to the fungal community (Rho = 0.218, *P* < 0.01), suggesting that the patterns of spatio-temporal variation were generally similar in both bacterial and fungal community distributions among tested plots and seasons.

**Fig. 4.**
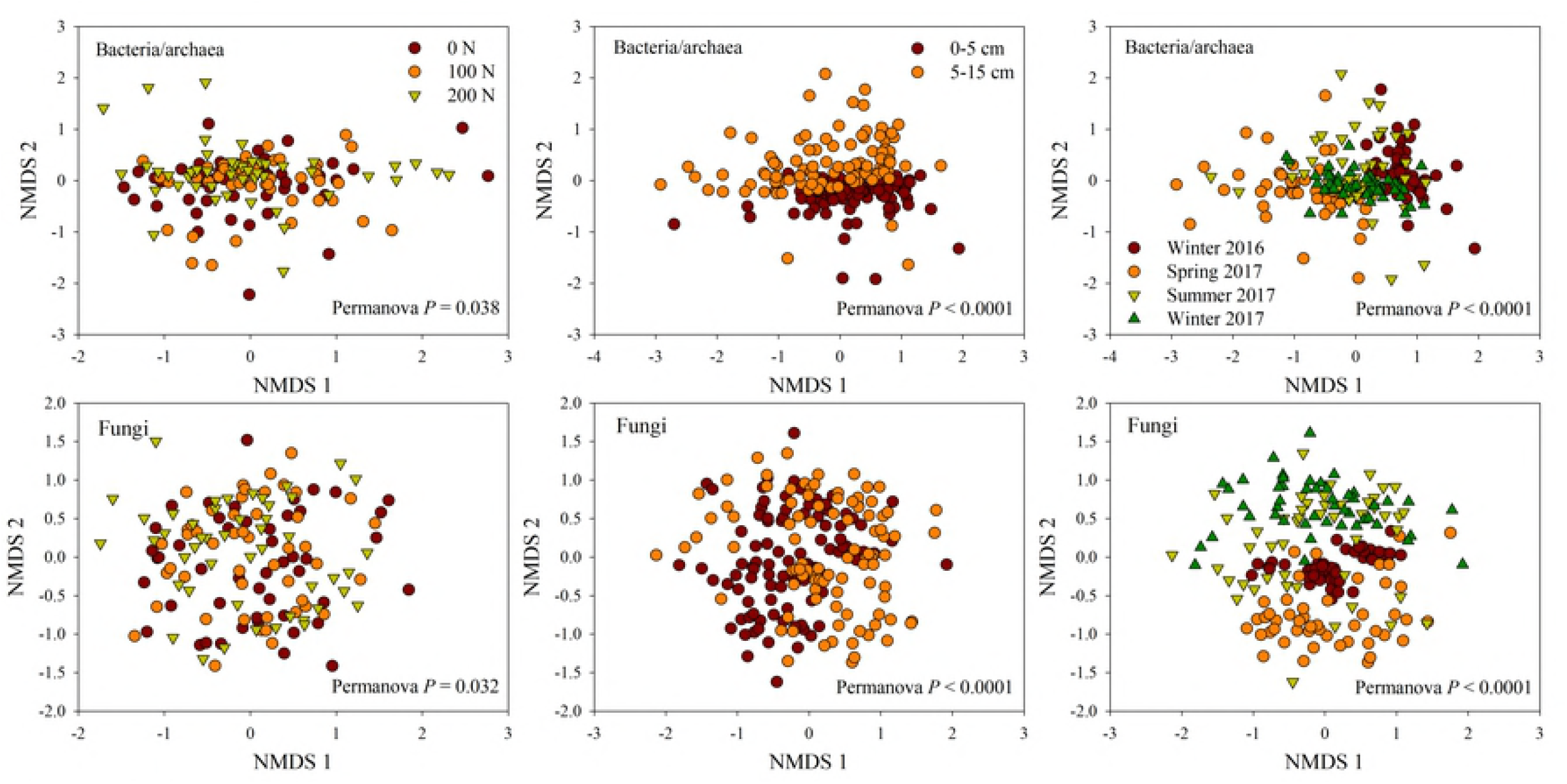
Non-metric multidimensional scaling (NMDS) analysis of bacterial/archaeal and fungal communities affected by three N fertilization levels (0, 100, and 200 kg N ha^-1^), two soil depth (0-5 and 5-15 cm), and four sampling seasons (Winter 2016, Spring, Summer, and Fall 2017). Permanova *P* values were also given.

**Table 1.**
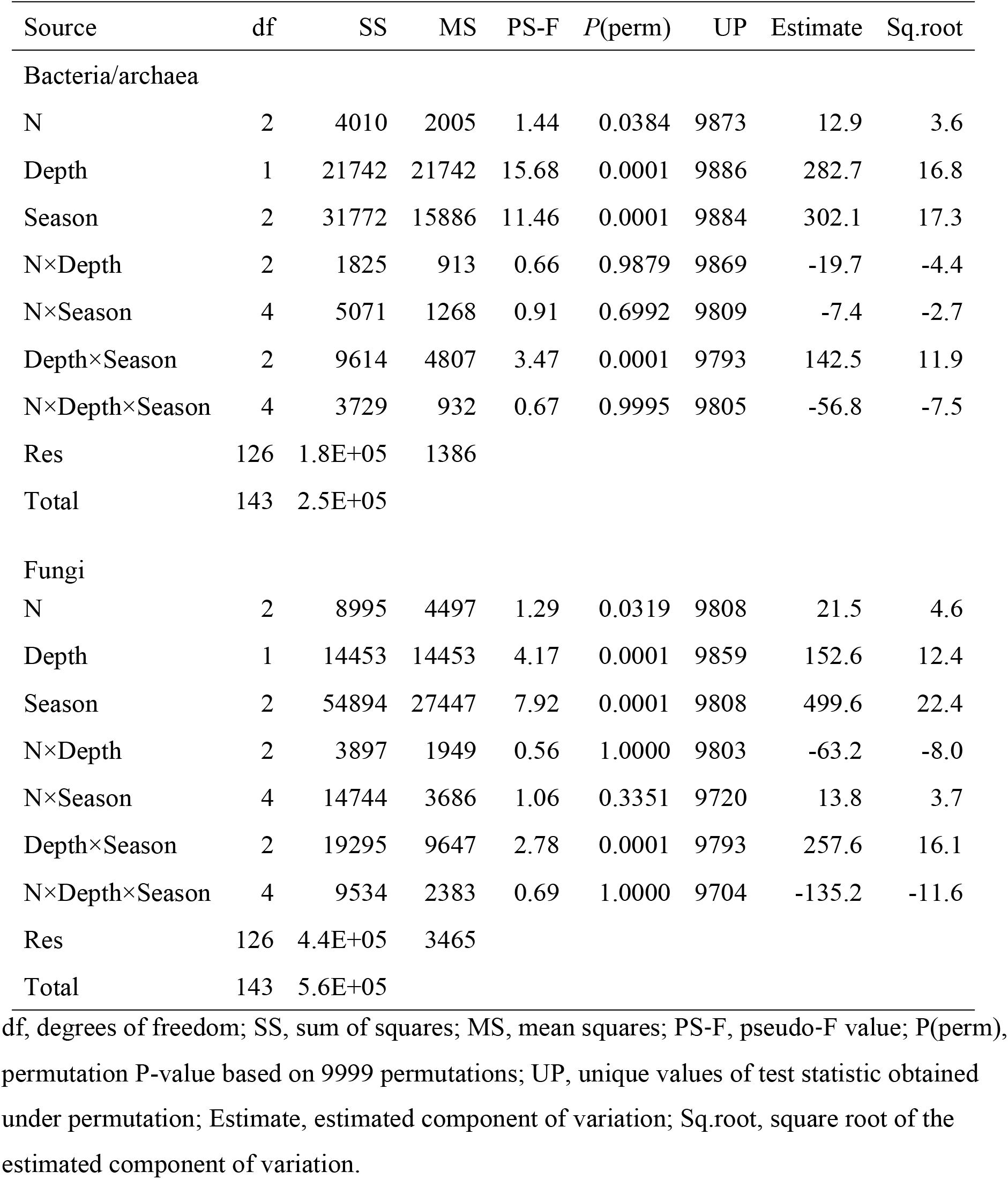
Three-factor Permanova results for differences in bacterial/archaeal and fungal community structure affected by three N fertilization levels (0, 100, and 200 kg N ha^-1^), two soil depths (0-5 and 5-15 cm), and three sampling seasons following N inputs (Spring, Summer, and Fall 2017). df, degrees of freedom; SS, sum of squares; MS, mean squares; PS-F, pseudo-F value; P(perm), permutation P-value based on 9999 permutations; UP, unique values of test statistic obtained under permutation; Estimate, estimated component of variation; Sq.root, square root of the estimated component of variation.

### Microbial taxonomic composition post-nitrogen addition

Because N level factors had no interaction with soil depth and sampling season (Table 1), N effects on microbial phylogenetic composition were assessed across both sampling depths and seasons (Fig. 5). Generally, N fertilization caused significant differences in the recovered genus level composition for prominent members of the bacterial/archaeal (6%) and the fungal (5%) communities, respectively (relative abundance > 0.01%) (Fig. 5). Specifically, for bacterial/archaeal community composition, N input at 200 kg N ha^-1^ significantly reduced the relative abundance of *Salinibacterium* and *Pseudonocardia* (Actinobacteria), *Caldilinear* (Chloroflexi), and *Desulfobulbus* (Proteobacteria), but increased *Sorangium* (Proteobacteria) (P < 0.05), indicating that these taxonomic groups were significantly altered by the synthetic N fertilizers. In the fungal community profiles, application of N fertilizers at 100 or 200 kg N ha^-1^ significantly decreased the proportion of *Archaeorhizomyces* (Ascomycota), as well as *Crepidotus* and *Uthatobasidium* (Basidiomycota) (P < 0.05).

**Fig. 5.**
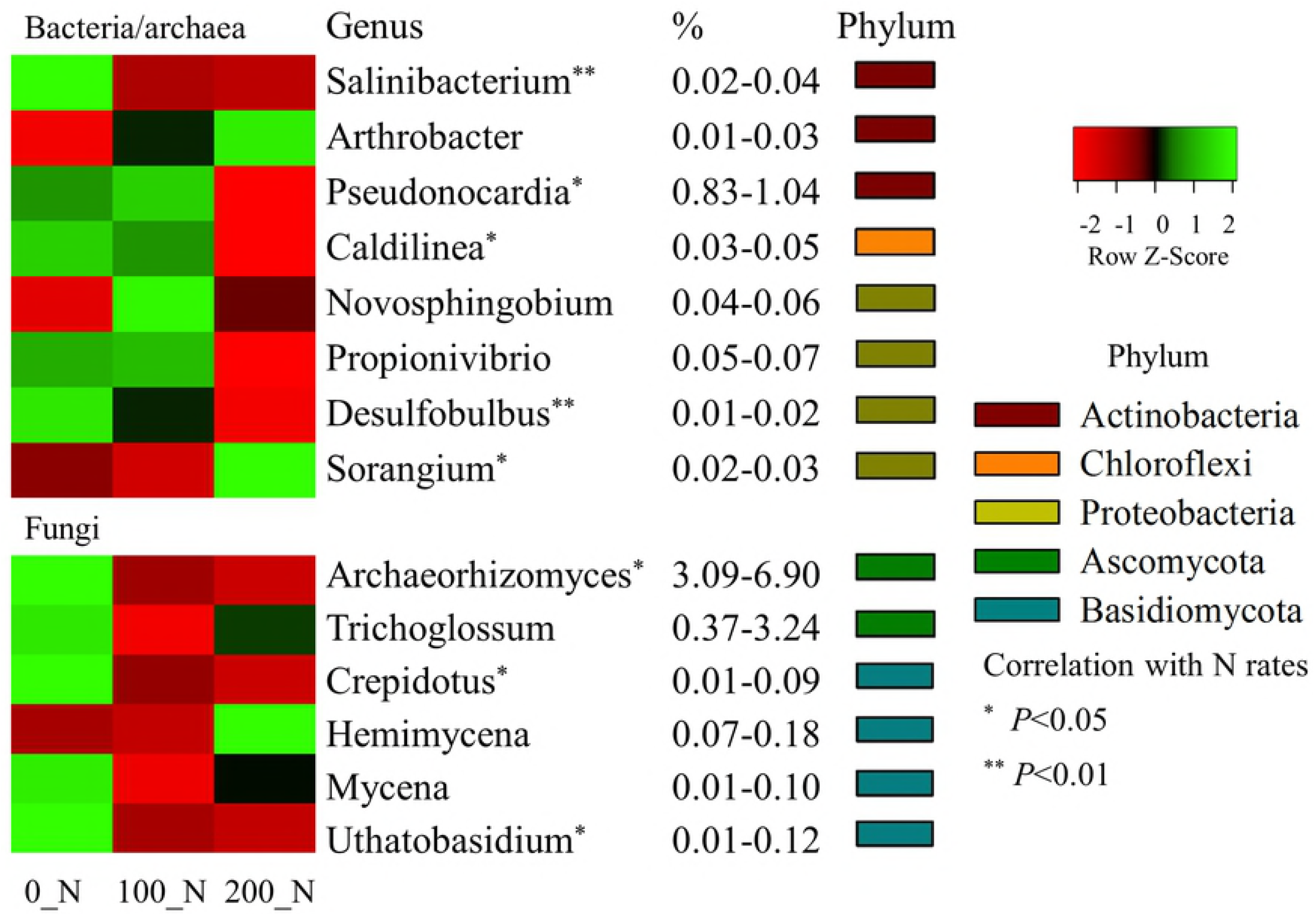
A heat map of relative abundance of bacterial/archaeal and fungal dominant taxonomic groups at genus level (average abundances > 0.1%) that were significantly affected by three N fertilization levels (0, 100, and 200 kg N ha^-1^). Asterisks indicate significant Spearman correlations of taxonomic abundance with N fertilization levels at α = 0.05.

Both soil depth and sampling season resulted in more significant alteration to bacterial/archaeal community composition than N application (Fig. 6). For example, 81% bacterial taxonomic groups at the genus level (with relative abundance > 0.1%) differed significantly between 0-5 and 5-15 cm of soil layers (P < 0.05), and significant variation occurred even at the phylum level. Generally, the 0-5 cm soil layer had a greater abundance of the phyla of Bacteroidetes, Planctomycetes, and Verrucomicrobia, whereas the phyla Chloroflexi, Nitrospirae, and Proteobacteria dominated the 5-15 cm soil layer (P < 0.05) (Fig. 6). Sampling season also caused significant changes in bacterial community composition with ~80% of bacterial genera significantly affected (P < 0.05) (Fig. 6), mostly in the prominent phyla of Acidobacteria, Actinobacteria, Bacteroidetes, and Verrucomicrobia, suggesting that these taxonomic groups were most responsive to temporal changes.

**Fig. 6.**
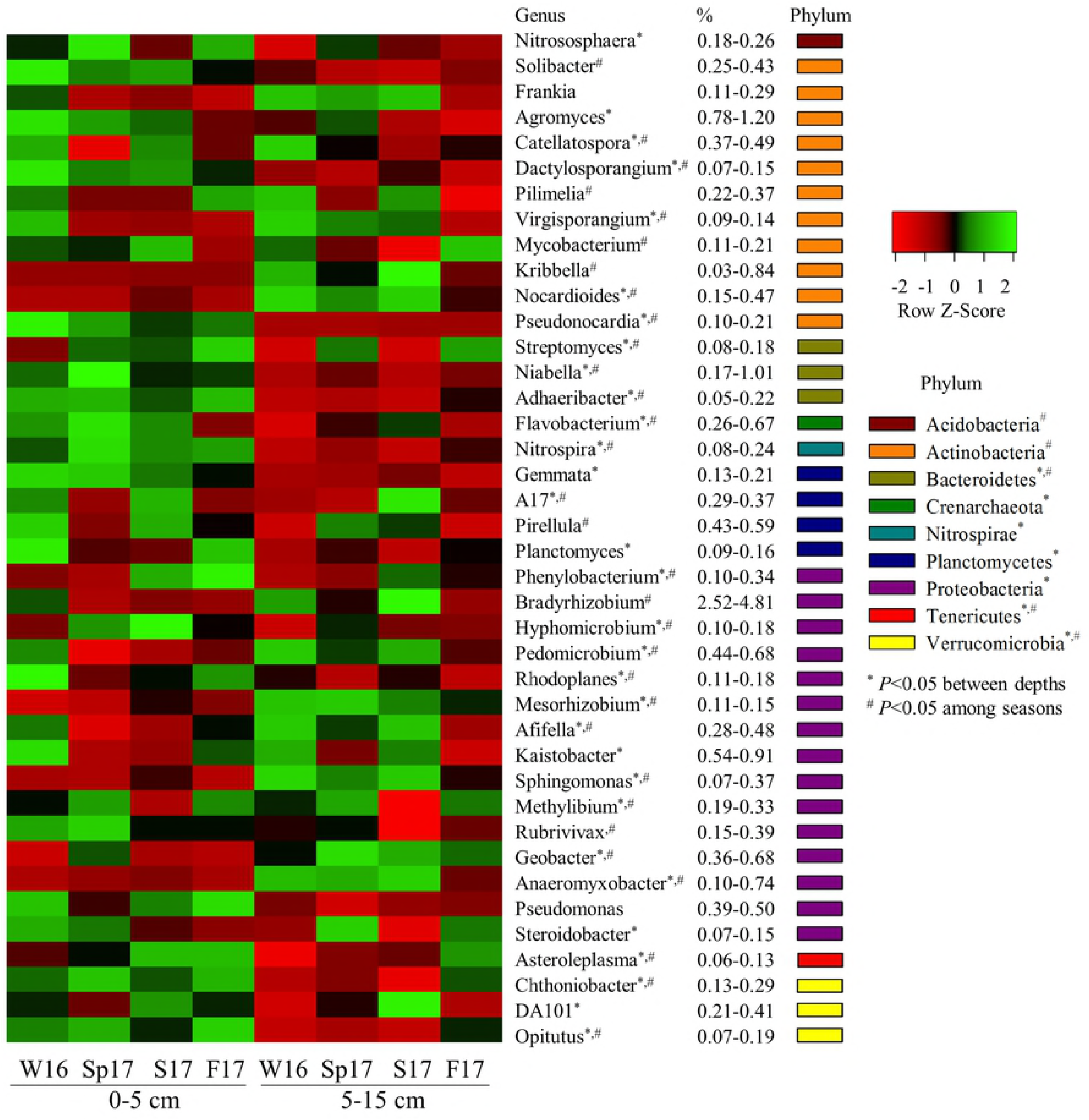
A heat map of relative abundance of bacterial/archaeal dominant taxonomic groups at genus level (average abundances > 0.1%) that were significantly variable between two soil depths (0-5 and 5-15 cm) and over four sampling seasons (Winter 2016, Spring, Summer, and Fall 2017). Asterisks indicate significant difference between two soil depths (0-5 and 5-15 cm) at α = 0.05. Number signs indicate significant difference over four sampling seasons (Winter 2016, Spring, Summer, and Fall 2017) at α = 0.05.

In the fungal community, only 54% prominent genera (of relative abundance >0.1%) showed a significant changes between two soil depths, in which members of the phyla of Ascomycota, Chytridiomycota, and Glomeromycota were more prevalent in top soil layer of 0-5 cm (P < 0.05) (Fig. 7). Approximately 90% of the prominent fungal taxonomic groups classified at the genus level (relative abundance > 0.1%) significantly varied over sampling seasons (P < 0. 05) (Fig. 7).

**Fig. 7.**
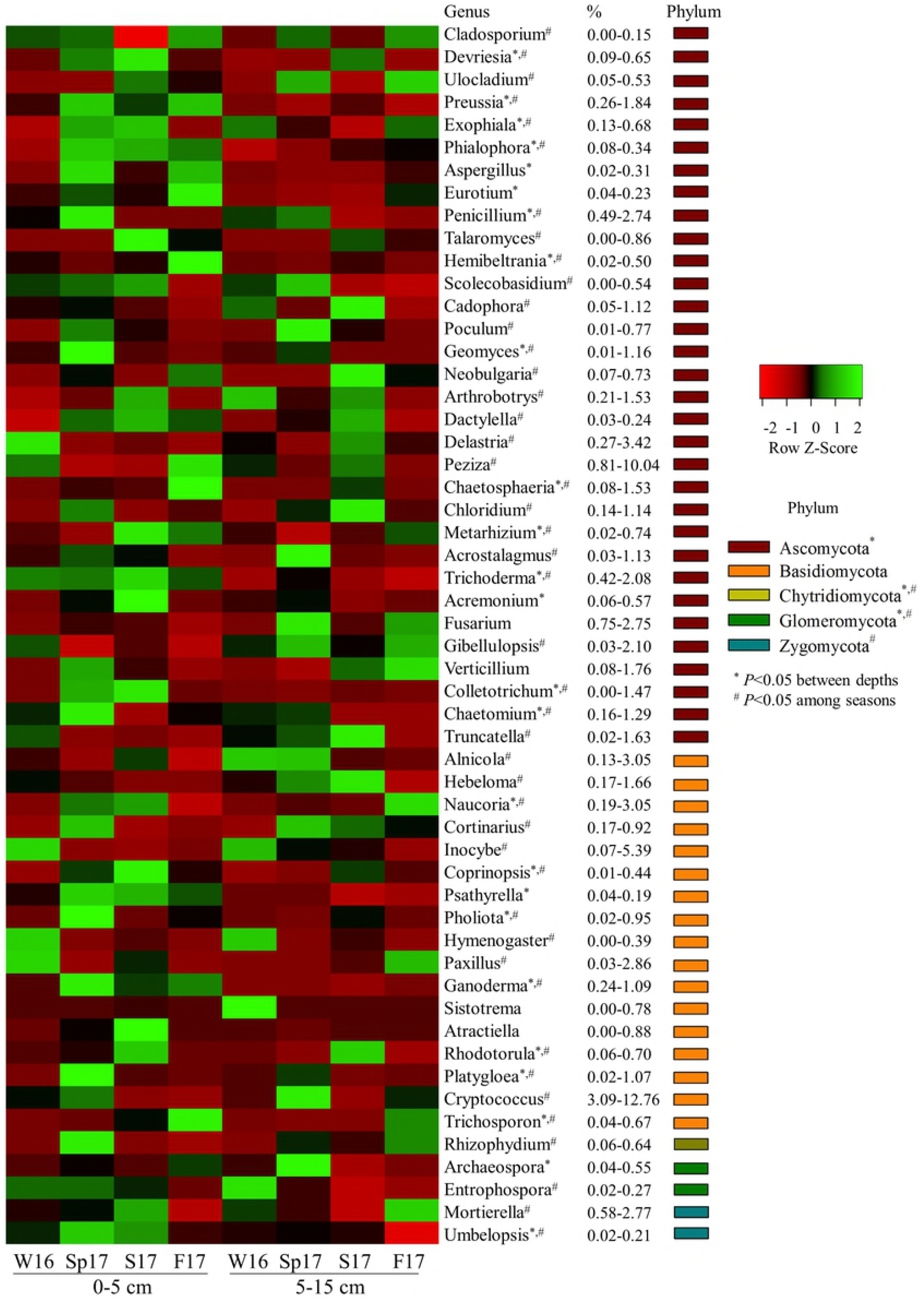
A heat map of relative abundance of fungal dominant taxonomic groups at genus level (average abundances > 0.1%) that were significantly variable between two soil depths (0-5 and 5-15 cm) and over four sampling seasons (Winter 2016, Spring, Summer, and Fall 2017). Asterisks indicate significant difference between two soil depths (0-5 and 5-15 cm) at α = 0.05. Number signs indicate significant difference over four sampling seasons (Winter 2016, Spring, Summer, and Fall 2017) at α = 0.05.

Venn diagrams were used to better visualize these changes of bacterial/archaeal and fungal taxonomic groups affected by soil depth and sampling season (Fig. 8). In bacteria, there were 61% of significantly affected genera shared by two factors of soil depth and sampling season, showing that most bacterial groups that differed between depths also responded to temporal change. In fungi, many more fungal taxonomic groups significantly varied across the four seasonal samples than depth difference (Fig. 8), indicating that temporal variation affected fungal community composition more significantly than spatial variation.

**Fig. 8.**
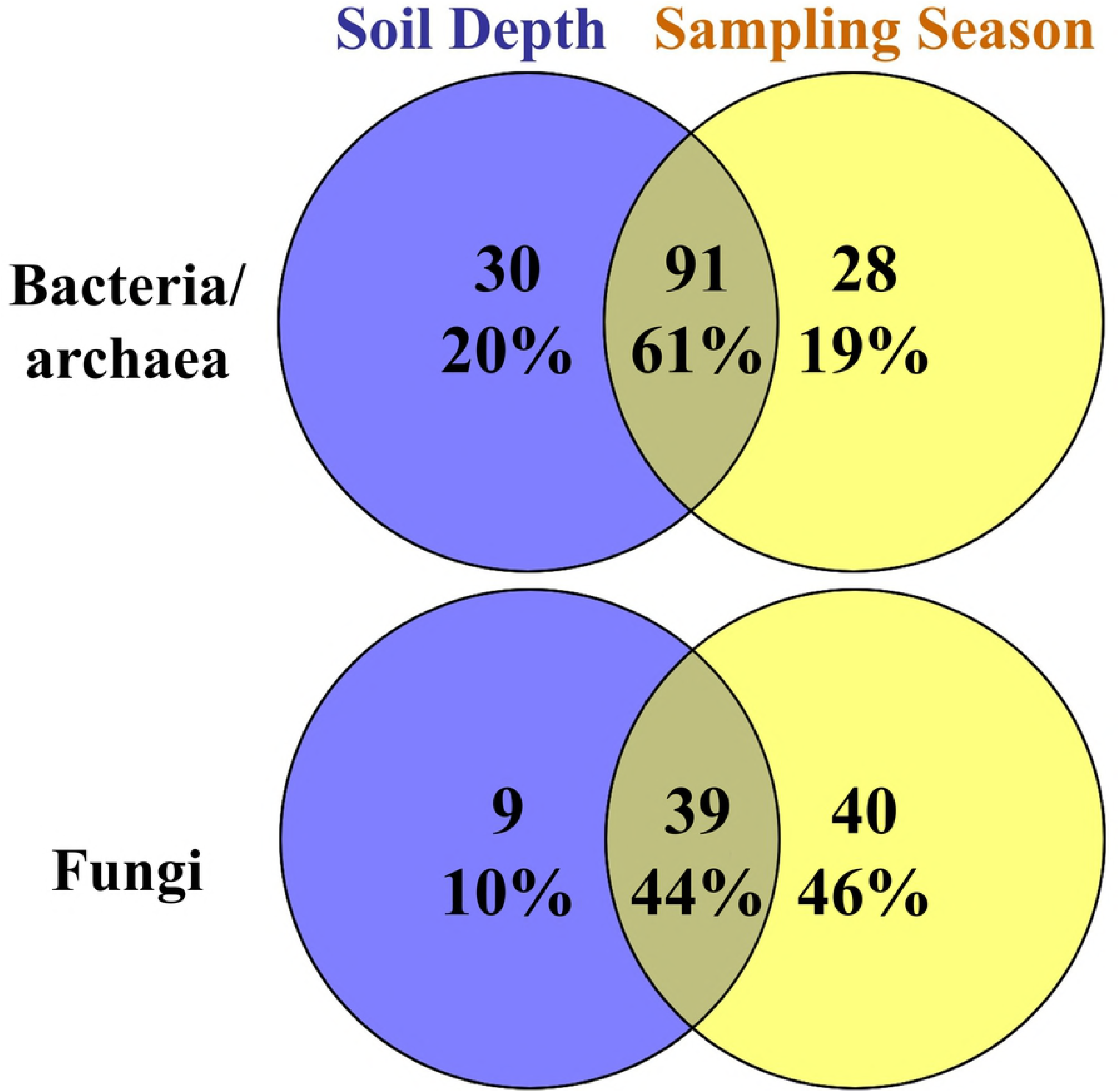
Venn’s diagrams showing significantly affected bacterial/archaeal and fungal dominant taxonomic groups at genus level (average abundances > 0.1%) shared between the factors of soil depth and sampling season.

### Microbial community associations with switchgrass yields and plant C/N contents

Compared to the control plots, N fertilization at 100 and 200 kg N ha^-1^ increased switchgrass yields by 43% and 171%, respectively (Table 2). In addition, N inputs also significantly increased plant N, but reduced relative C content and C/N ratios (P < 0.05) as measured at the end of the growing season. The DistLM analysis showed that switchgrass yields were significantly correlated with the community structure of bacteria/archaea and fungi, but explained only a small portion of variation, *i.e.,* 2.6%, 1.2%, in bacterial/archaeal and fungal profiles, respectively (P < 0.01) (Table 2), suggesting a small but significant correlation between above-ground switchgrass growth and below-ground microbiomes through N fertilization.

**Table 2.** Above-ground biomass yields and plant C/N contents of switchgrass affected by N fertilization levels as well as their association with community structure of bacteria/archaea and fungi by marginal test of DistLM. Different letters within each column indicate significant effects by N fertilization levels at α=0.05. The *, **, and *** indicate significant DistLM relationship at α=0.05, 0.01, and 0.001, respectively.

|  | Yield (Mg Ha<sup>-1</sup>) | C (%) | N (%) | C/N |
| --- | --- | --- | --- | --- |
| N level (kg N ha<sup>-1</sup>) |  |  |  |  |
| 0 | 29.6 b | 50.43 a | 0.28 a | 189.46 a |
| 100 | 42.4 ab | 49.70 b | 0.83 b | 74.73 b |
| 200 | 80.1 a | 49.47 b | 0.98 b | 60.62 b |
| Proportion of explained variation |  |  |  |  |
| Bacteria/archaea | 2.6%*** | 1.2%* | 1.0% | 0.6% |
| Fungi | 1.2%** | 1.0% | 1.0% | 0.8% |

## Discussion

### Short-term N effects on microbial communities

Long-term N input can alter microbial composition and diversity, mainly due to N-induced soil acidification and fertility decline [10]. Many long-term studies have reported that N fertilization not only reduces below-ground biodiversity but also shifts bacterial composition at the phylum level, for groups such as Proteobacteria, Acidobacteria and Actinobacteria [44–48]. Field studies focusing on short-term effects of N fertilization on microbial communities however are limited in number for comparison. In our study, one-time fertilization did not affect the richness and diversity of soil microbial communities, but caused structural changes in both bacterial/archaeal and fungal community composition (Fig. 4). Our work suggests that some phylogenetic groups of bacteria and fungi might quickly react to N inputs, even when soil properties are not significantly modified by short-term N fertilization. These N effects were consistent across two soil depths and four sampling seasons because there was no significant interaction between N and depth/season (Table 1). We also observed that the one-time N amendment appeared to directly repress some bacterial and fungal groups based on the negative relationship of relative abundance with N levels, for example bacterial genus *Pseudonocardia*, and fungal genus *Archaeorhizomyces* (Fig. 5).

*Pseudonocardia* is a common endophytic Actinomycete frequently isolated from host plant tissues [49], which has been reported to achieve associative nitrogen fixation without the formation of nodules [50] and protect their hosts against soil-borne pathogenic infection through producing antibiotics or siderophores [51, 52]. As a free living diazotrophic Actinomycete, it has also been reported to be prominent in nutrient limited environments [53, 54] or low-input agroecosystems [55, 56], due to its low requirement for N. Based on sequencing of 16S rRNA genes, it was also found that *Pseudonocardia* OTUs were reduced in the fertilized plant rhizosphere of Canola *(Brassica napus)* [57]. Our results support that the relative abundances of *Pseudonocardia* are significantly and negatively associated with N fertilization (Fig. 5), suggesting that even short-term N inputs might acutely suppress this associative nitrogen fixer in switchgrass cultivated lands.

The *Archaeorhizomyces* are an ancient class of ubiquitous soil fungi [58], which are neither mycorrhizal nor pathogenic, but may be root endophytic or free-living saprophytes [59]. This group was first discovered in tundra soils [60] using rRNA-based sequencing, but was only isolated into culture more recently [58] and very little is definitively known about the physiology and ecology of this group of organisms and this knowledge comes only from only one extant isolate of the broad class of organisms. By investigating how organic matter accumulation and forest fertility influences fungal community composition, it was found from ITS rRNA gene analyses that *Archaeorhizomyces* dominated root-associated Ascomycetes and there abundance significantly correlated with a fertility gradient in European boreal forests [61]. Moreover, it has been shown that the relative abundance of *Archaeorhizomyces* in grasslands is greatly stimulated by amendment of the biofertilizers *Trichoderma* [62] and correlations between soil properties and fungal abundance suggested that soil P availability (rather than N) may be a controlling factor for *Archaeorhizomyces* relative abundance. However, in our study, inorganic N fertilization significantly reduced the relative abundance of *Archaeorhizomyces*, which was one of the dominant groups of the in Ascomycota present in our study at 3.1-6.9% relative abundance (Fig. 5). Further studies on the ecology of these diverse fungi are clearly needed through both additional rRNA gene amplicon studies in natural systems, as well as the isolation of additional representatives for ecophysiological analyses.

### Spatial heterogeneity in microbial communities

Several studies have reported that soil bacterial and fungal diversity levels can either decrease [23, 63, 64], remain unchanged [25, 65, 66] or increase [67] with soil depth. We consistently observed reduced community richness and diversity in 5-15 cm compared to 0-5 cm soil layers for both bacterial/archaeal and fungal communities (Fig. 3). Since plant residue serves as a key carbon source for soil microbes, the vertical distribution of microbial communities is likely to reflect the different available organic matter content with soil depths for microbial decomposers [64]. For example, the surface soil may have more easily decomposable carbon directly derived from crop residues, with more diverse groups of microbes able to access the labile organic materials in this niche [68] whereas subsurface soils may harbor relatively more recalcitrant carbon sources or be more dependent on root inputs. We also observed less soil C and N in 5-15 cm soil layers, further suggesting nutrient levels may be among the factors driving these depth related patterns in diversity.

Compared to the small amount of community variation attributable to N addition, we observed more significant shifts in both bacterial/archaeal and fungal community composition between soil depths. Generally, Bacteroidetes, Planctomycetes, and Verrucomicrobia were more abundant in the 0-5 cm soil layer. This spatial differentiation of the dominant bacterial groups by soil depth was consistent to many previous studies. For example, it has been shown that bacterial community composition was significantly altered at different soil depths, which was associated primarily with a decline of Bacteroidetes with depth [23]. Others have also reported that Verrucomicrobia exhibit higher relative abundance in the surface soils [25, 67]. In contrast, our results showed that the 5-15 cm soil layer had greater abundance in the phyla of Chloroflexi, Nitrospirae, and Proteobacteria. Similarly, it is also demonstrated that as soil depth increased, the relative abundance of Proteobacteria increased and it became the dominant bacterial group in subsoil [65]. Though the overall Proteobacteria were more abundant in 5-15 cm soils, the class Betaproteobacteria was most abundant in 0-5 cm, which was also found in other study [69]. Similar to our results, others have also reported that Chloroflexi [66, 67] and Nitrospirae [70] increase in abundance with soil depth.

In this study, we observed that fungal community also showed strong vertical distribution patterns in the major groups, such as Ascomycota, Chytridiomycota, and Glomeromycota, which were more abundant in top 0-5 cm soil layer; however, compared to bacterial community, there were overall fewer fungal taxonomic groups that differed between soil depths (Fig. 6 and 7). Several studies have highlighted the ecological significance of vertically distinct fungal communities. For example, using pyrosequencing of ITS amplicon, others have found a decrease in relative abundance of Ascomycota with increasing soil depth, whereas Zygomycota showed the opposite trend [63]. At finer taxonomic scales, it was reported that Sordariomycetes of the phylum Ascomycota decrease with soil depth [70], and this pattern was similar in our study. Others have also shown overall fungal communities pattern were highly variable with soil depth, where deeper soil have some distinct fungal groups, but significantly less overall diversity [64] similar to what we observe here.

### Temporal variation in microbial communities

We found significant temporal changes in alpha diversity in both bacterial and fungal communities (Fig. 3). Similarly, it has been shown that bacterial community alpha diversity varied more substantially than beta diversity over time, and in this case exceeded the variability between land-use types [24]. Thus, it was suggested that temporal differences in rhizodeposition may be a controlling factor to affect soil bacterial diversity. Our data show that seasonal variation is found in most dominant phylogenetic groups of bacteria, such as Acidobacteria, Actinobacteria, Bacteroidetes, Chloroflexi, and Verrucomicrobia (Fig. 6). Interestingly Acidobacteria, Bacteroidetes, Betaproteobacteria, Deltaproteobacteria, Gammaproteobacteria, and Verrucomicrobia all had similar seasonal patterns, which were opposite to those of Actinobacteria, Chloroflexi, and Alphaproteobacteria. That dominant bacterial phyla, such as Actinobacteria and Betaproteobacteria, shift with seasonally to temporal patterns has been reported previously [65]. In addition, it has been shown that seasonal dynamics often appear to be coherent within taxonomic lineages, in which Acidobacteria and Proteobacteria are more prevalent in summer, whereas Actinobacteria and Chloroflexi increase in winter [71].

In our experiment, the fungal phyla Chytridiomycota, Glomeromycota, and Zygomycota, were also found to vary significantly over the different seasonal sampling times (Fig. 7). Similarly, it was reported that the number of fungal species belonging to Ascomycota and Glomeromycota increase in summer, whereas Basidiomycota were dominant in winter [72] or that Ascomycota, Basidiomycota, and Zygomycota are variable from spring to winter [73]. It has been suggested that changes in litter decomposition and phytosynthate allocation contribute to the seasonal variations of fungal community [74] as well as the direct effects of soil moisture and temperature [24]. However, contrary to this we did not observe distinct seasonal changes in the overall dominance patterns of Ascomycota or Basidiomycota in our study.

## Conclusions

With the aid of high-throughput 16S rRNA gene and ITS region amplicon sequencing, we found highly diverse and dynamic communities across this 8 year-old switchgrass field. The one-time application of N fertilization significantly stimulated switchgrass growth and N uptake, and subtly but significantly shifted below-ground bacterial and fungal communities, with the bacterial genus *Pseudonocardia* and *Archaeorhizomyces* fungi negatively responsive to N inputs. However, these shifts took place within the context of much larger spatial and temporal variation in the microbial community. These large spatial and seasonal fluctuations in microbial communities reinforce the importance of robust sampling designs and should caution against overinterpretation of studies based on one-time sampling events. Further studies should aim at studying the ecological and physiological mechanisms of responses to N fertilization by these microbes and how these may influence ecosystem functions.

## Data Availability Statement

The 16S rRNA gene and ITS region amplicon sequences were deposited on NCBI Sequence Read Archive (SRA) database under the BioProject accession number of PRJNA512218. Soil metagenome sequences were uploaded to Rapid Annotation using Subsystems Technology for Metagenomes (MG-RAST; http://metagenomics.anl.gov) under project accession number mgp22000.

## Author Contributions

Conceived and designed the experiments: HC CWS. Performed the experiments: HC ZKY DY RHM SJL DMK DH CWS. Analyzed the data: HC CWS. Contributed to the writing of the manuscript: HC MAC RLH GW FEL CWS.

## Supporting Information

**Fig. S1.** Relative abundances of soil metagenomes annotated in RefSeq database (RefSeq), and functional classification annotated in SEED Subsystems database (SEED Subsystems) in composite soils sampled at two soil depths (0-5 and 5-15 cm) in Winter 2016. Asterisks indicate significant difference at α = 0.05 between two soil depths.

**Fig. S2.** Non-metric multidimensional scaling (NMDS) analysis of 24 putative functions at KEGG level 1 predicted by PICRUSt based on 16S rRNA amplicons compared with 4 soil metagenomes annotated in KO database (KEGG level 1) using soils sampled in Winter 2016. Permanova *P* values were also given.

## References

1. Demirbas A. Political, economic and environmental impacts of biofuels: A review. Applied Energy. 2009;86, Supplement 1(0):S108–S17.

2. Jagger A. Biofuels for transport in 2050. Biofuels, Bioproducts and Biorefining. 2011;5(5):481–5.

3. Wang Z, Smyth TJ, Crozier CR, Gehl RJ, Heitman AJ. Yield and Nitrogen Removal of Bioenergy Grasses as Influenced by Nitrogen Rate and Harvest Management in the Coastal Plain Region of North Carolina. BioEnergy Research. 2018;11(1):44–53.

4. Fike JH, Pease JW, Owens VN, Farris RL, Hansen JL, Heaton EA, et al. Switchgrass nitrogen response and estimated production costs on diverse sites. Gcb Bioenergy. 2017;9(10):1526–42.

5. Gopalakrishnan G, Cristina Negri M, Snyder SW. A novel framework to classify marginal land for sustainable biomass feedstock production. Journal of environmental quality. 2011;40(5):1593–600.

6. Toma YO, FernÁNdez FG, Sato S, Izumi M, Hatano R, Yamada T, et al. Carbon budget and methane and nitrous oxide emissions over the growing season in a Miscanthus sinensis grassland in Tomakomai, Hokkaido, Japan. GCB Bioenergy. 2011;3(2):116–34.

7. Nazir R, Warmink JA, Boersma H, Van Elsas JD. Mechanisms that promote bacterial fitness in fungal-affected soil microhabitats. FEMS Microbiology Ecology. 2010;71(2):169–85.

8. Nielsen U, Ayres E, Wall D, Bardgett R. Soil biodiversity and carbon cycling: a review and synthesis of studies examining diversity–function relationships. European Journal of Soil Science. 2011;62(1):105–16.

9. Van Der Heijden MG, Bardgett RD, Van Straalen NM. The unseen majority: soil microbes as drivers of plant diversity and productivity in terrestrial ecosystems. Ecology letters. 2008; 11(3):296–310.

10. Dai Z, Su W, Chen H, Barberán A, Zhao H, Yu M, et al. Long-term nitrogen fertilization decreases bacterial diversity and favors the growth of Actinobacteria and Proteobacteria in agro-ecosystems across the globe. Global change biology. 2018.

11. Ramirez KS, Craine JM, Fierer N. Consistent effects of nitrogen amendments on soil microbial communities and processes across biomes. Global Change Biology. 2012;18(6):1918–27.

12. Leff JW, Jones SE, Prober SM, Barberán A, Borer ET, Firn JL, et al. Consistent responses of soil microbial communities to elevated nutrient inputs in grasslands across the globe. Proceedings of the National Academy of Sciences. 2015;112(35):10967–72.

13. Rousk J, Bååth E, Brookes PC, Lauber CL, Lozupone C, Caporaso JG, et al. Soil bacterial and fungal communities across a pH gradient in an arable soil. The ISME journal. 2010;4(10):1340.

14. Bardgett RD, Lovell RD, Hobbs PJ, Jarvis SC. Seasonal changes in soil microbial communities along a fertility gradient of temperate grasslands. Soil Biology and Biochemistry. 1999;31(7):1021–30.

15. Tlustos P, Willison T, Baker J, Murphy D, Pavlikova D, Goulding K, et al. Short-term effects of nitrogen on methane oxidation in soils. Biology and Fertility of Soils. 1998;28(1):64–70.

16. Wang Q, Wang S, Liu Y. Responses to N and P fertilization in a young Eucalyptus dunnii plantation: microbial properties, enzyme activities and dissolved organic matter. Applied Soil Ecology. 2008;40(3):484–90.

17. Wang C, Long R, Wang Q, Liu W, Jing Z, Zhang L. Fertilization and litter effects on the functional group biomass, species diversity of plants, microbial biomass, and enzyme activity of two alpine meadow communities. Plant and Soil. 2010;331(–2):377–89.

18. Jang I, Lee S, Zoh K-D, Kang H. Methane concentrations and methanotrophic community structure influence the response of soil methane oxidation to nitrogen content in a temperate forest. Soil Biology and Biochemistry. 2011;43(3):620–7.

19. Deslippe JR, Egger KN, Henry GH. Impacts of warming and fertilization on nitrogen-fixing microbial communities in the Canadian High Arctic. FEMS microbiology ecology. 2005;53(1):41–50.

20. Piceno Y, Lovell C. Stability in natural bacterial communities: I. Nutrient addition effects on rhizosphere diazotroph assemblage composition. Microbial ecology. 2000;39(1):32–40.

21. Karp A, Shield I. Bioenergy from plants and the sustainable yield challenge. New Phytologist. 2008;179(1):15–32.

22. McLaughlin SB, Kszos LA. Development of switchgrass (Panicum virgatum) as a bioenergy feedstock in the United States. Biomass and Bioenergy. 2005;28(6):515–35.

23. Eilers KG, Debenport S, Anderson S, Fierer N. Digging deeper to find unique microbial communities: the strong effect of depth on the structure of bacterial and archaeal communities in soil. Soil Biology and Biochemistry. 2012;50:58–65.

24. Lauber CL, Ramirez KS, Aanderud Z, Lennon J, Fierer N. Temporal variability in soil microbial communities across land-use types. The ISME journal. 2013;7(8):1641.

25. Bai R, Wang J-T, Deng Y, He J-Z, Feng K, Zhang L-M. Microbial community and functional structure significantly varied among distinct types of paddy soils but responded differently along gradients of soil depth layers. Frontiers in microbiology. 2017;8:945.

26. Dong H-Y, Kong C-H, Wang P, Huang Q-L. Temporal variation of soil friedelin and microbial community under different land uses in a long-term agroecosystem. Soil Biology and Biochemistry. 2014;69:275–81.

27. Will C, Thürmer A, Wollherr A, Nacke H, Herold N, Schrumpf M, et al. Horizon-specific bacterial community composition of German grassland soils, as revealed by pyrosequencing-based analysis of 16S rRNA genes. Applied and environmental microbiology. 2010;76(20):6751–9.

28. Lewandowski TE, Forrester JA, Mladenoff DJ, Stoffel JL, Gower ST, D’Amato AW, et al. Soil microbial community response and recovery following group selection harvest: Temporal patterns from an experimental harvest in a US northern hardwood forest. Forest Ecology and Management. 2015;340:82–94.

29. Kramer S, Marhan S, Haslwimmer H, Ruess L, Kandeler E. Temporal variation in surface and subsoil abundance and function of the soil microbial community in an arable soil. Soil Biology and Biochemistry. 2013;61:76–85.

30. Cregger MA, Schadt CW, McDowell NG, Pockman WT, Classen AT. Response of the soil microbial community to changes in precipitation in a semiarid ecosystem. Applied and environmental Microbiology. 2012;78(24):8587–94.

31. Stevenson BA, Hunter DW, Rhodes PL. Temporal and seasonal change in microbial community structure of an undisturbed, disturbed, and carbon-amended pasture soil. Soil Biology and Biochemistry. 2014;75:175–85.

32. DeBruyn JM, Bevard DA, Essington ME, McKnight JY, Schaeffer SM, Baxter HL, et al. Field-grown transgenic switchgrass (Panicum virgatum L.) with altered lignin does not affect soil chemistry, microbiology, and carbon storage potential. Gcb Bioenergy. 2017;9(6):1100–9.

33. Kennedy RK. The sickledrat: a circular quadrat modification useful in grassland studies. Rangeland Ecology & Management/Journal of Range Management Archives. 1972;25(4):312–3.

34. Lundberg DS, Yourstone S, Mieczkowski P, Jones CD, Dangl JL. Practical innovations for high-throughput amplicon sequencing. Nature methods. 2013;10(10):999.

35. Cregger M, Veach A, Yang Z, Crouch M, Vilgalys R, Tuskan G, et al. The Populus holobiont: dissecting the effects of plant niches and genotype on the microbiome. Microbiome. 2018;6(1):31.

36. Martin M. Cutadapt removes adapter sequences from high-throughput sequencing reads. EMBnet journal. 2011;17(1):pp. 10–2.

37. Caporaso JG, Kuczynski J, Stombaugh J, Bittinger K, Bushman FD, Costello EK, et al. QIIME allows analysis of high-throughput community sequencing data. Nature Methods. 2010;7(5):335–6.

38. Wang Q, Garrity GM, Tiedje JM, Cole JR. Naive Bayesian classifier for rapid assignment of rRNA sequences into the new bacterial taxonomy. Applied and environmental microbiology. 2007;73(16):5261–7.

39. Langille MG, Zaneveld J, Caporaso JG, McDonald D, Knights D, Reyes JA, et al. Predictive functional profiling of microbial communities using 16S rRNA marker gene sequences. Nature biotechnology. 2013;31(9):814–21.

40. Meyer F, Paarmann D, D'Souza M, Olson R, Glass EM, Kubal M, et al. The metagenomics RAST server-a public resource for the automatic phylogenetic and functional analysis of metagenomes. BMC bioinformatics. 2008;9(1):386.

41. Anderson M. DISTLM v. 5: a FORTRAN computer program to calculate a distance-based multivariate analysis for a linear model. Department of Statistics, University of Auckland, New Zealand. 2004;10:2016.

42. Babicki S, Arndt D, Marcu A, Liang Y, Grant JR, Maciejewski A, et al. Heatmapper: web- enabled heat mapping for all. Nucleic acids research. 2016;44(W1):W147–W53.

43. Oliveros J. An interactive tool for comparing lists with Venn’s diagrams (2007-2015).

44. Zhou J, Guan D, Zhou B, Zhao B, Ma M, Qin J, et al. Influence of 34-years of fertilization on bacterial communities in an intensively cultivated black soil in northeast China. Soil Biology and Biochemistry. 2015;90:42–51.

45. Xun W, Zhao J, Xue C, Zhang G, Ran W, Wang B, et al. Significant alteration of soil bacterial communities and organic carbon decomposition by different long-term fertilization management conditions of extremely low-productivity arable soil in South China. Environmental microbiology. 2016;18(6):1907–17.

46. Ling N, Chen D, Guo H, Wei J, Bai Y, Shen Q, et al. Differential responses of soil bacterial communities to long-term N and P inputs in a semi-arid steppe. Geoderma. 2017;292:25–33.

47. Coolon JD, Jones KL, Todd TC, Blair JM, Herman MA. Long-term nitrogen amendment alters the diversity and assemblage of soil bacterial communities in tallgrass prairie. PLoS One. 2013;8(6):e67884.

48. Pan Y, Cassman N, de Hollander M, Mendes LW, Korevaar H, Geerts RH, et al. Impact of long-term N, P, K and NPK fertilization on the composition and potential functions of the bacterial community in grassland soil. FEMS microbiology ecology. 2014;90(1):195–205.

49. Hamedi J, Mohammadipanah F. Biotechnological application and taxonomical distribution of plant growth promoting actinobacteria. Journal of industrial microbiology & biotechnology. 2015;42(2):157–71.

50. Valdés M, Pérez N-O, Estrada-de Los Santos P, Caballero-Mellado J, Pena-Cabriales JJ, Normand P, et al. Non-Frankia actinomycetes isolated from surface-sterilized roots of Casuarina equisetifolia fix nitrogen. Applied and environmental microbiology. 2005;71(1):460–6.

51. Janso JE, Carter GT. Biosynthetic potential of phylogenetically unique endophytic actinomycetes from tropical plants. Applied and environmental microbiology. 2010;76(13):4377–86.

52. Wang R, Zhang H, Sun L, Qi G, Chen S, Zhao X. Microbial community composition is related to soil biological and chemical properties and bacterial wilt outbreak. Scientific Reports. 2017;7(1):343.

53. Zhang Y-Q, Liu H-Y, Chen J, Yuan L-J, Sun W, Zhang L-X, et al. Diversity of culturable actinobacteria from Qinghai–Tibet plateau, China. Antonie Van Leeuwenhoek. 2010;98(2):213–23.

54. De Mandal S, Chatterjee R, Kumar NS. Dominant bacterial phyla in caves and their predicted functional roles in C and N cycle. BMC microbiology. 2017;17(1):90.

55. Yin C, Mueth N, Hulbert S, Schlatter D, Paulitz TC, Schroeder K, et al. Bacterial communities on wheat grown under long-term conventional tillage and no-till in the Pacific Northwest of the United States. Phytobiomes. 2017;1(2):83–90.

56. Dorr de Quadros P, Zhalnina K, Davis-Richardson A, Fagen JR, Drew J, Bayer C, et al. The effect of tillage system and crop rotation on soil microbial diversity and composition in a subtropical acrisol. Diversity. 2012;4(4):375–95.

57. Monreal C, Zhang J, Koziel S, Vidmar J, González M, Matus F, et al. Bacterial community structure associated with the addition of nitrogen and the dynamics of soluble carbon in the rhizosphere of canola (Brassica napus) grown in a Podzol. Rhizosphere. 2018;5:16–25.

58. Rosling A, Cox F, Cruz-Martinez K, Ihrmark K, Grelet G-A, Lindahl BD, et al. Archaeorhizomycetes: unearthing an ancient class of ubiquitous soil fungi. Science. 2011;333(6044):876–9.

59. Rosling A, Timling I, Taylor DL. Archaeorhizomycetes: Patterns of distribution and abundance in soil. Genomics of Soil-and Plant-Associated Fungi: Springer; 2013. p. 333–49.

60. Schadt CW, Martin AP, Lipson DA, Schmidt SK. Seasonal dynamics of previously unknown fungal lineages in tundra soils. Science. 2003;301(5638):1359–61.

61. Kyaschenko J, Clemmensen KE, Karltun E, Lindahl BD. Below-ground organic matter accumulation along a boreal forest fertility gradient relates to guild interaction within fungal communities. Ecology letters. 2017;20(12):1546–55.

62. Zhang F, Huo Y, Cobb AB, Luo G, Zhou J, Yang G, et al. Trichoderma biofertilizer links to altered soil chemistry, altered microbial communities, and improved grassland biomass. Frontiers in microbiology. 2018;9.

63. Santalahti M, Sun H, Jumpponen A, Pennanen T, Heinonsalo J. Vertical and seasonal dynamics of fungal communities in boreal Scots pine forest soil. FEMS microbiology ecology. 2016;92(11):fiw170.

64. Schlatter DC, Kahl K, Carlson B, Huggins DR, Paulitz T. Fungal community composition and diversity vary with soil depth and landscape position in a no-till wheat-based cropping system. FEMS microbiology ecology. 2018.

65. Bevivino A, Paganin P, Bacci G, Florio A, Pellicer MS, Papaleo MC, et al. Soil bacterial community response to differences in agricultural management along with seasonal changes in a Mediterranean region. PLoS One. 2014;9(8):e105515.

66. Steven B, Gallegos-Graves LV, Belnap J, Kuske CR. Dryland soil microbial communities display spatial biogeographic patterns associated with soil depth and soil parent material. FEMS microbiology ecology. 2013;86(1):101–13.

67. Mueller RC, Belnap J, Kuske CR. Soil bacterial and fungal community responses to nitrogen addition across soil depth and microhabitat in an arid shrubland. Frontiers in microbiology. 2015;6:891.

68. Boer Wd, Folman LB, Summerbell RC, Boddy L. Living in a fungal world: impact of fungi on soil bacterial niche development. FEMS microbiology reviews. 2005;29(4):795–811.

69. Lüdemann H, Arth I, Liesack W. Spatial changes in the bacterial community structure along a vertical oxygen gradient in flooded paddy soil cores. Applied and Environmental Microbiology. 2000;66(2):754–62.

70. Ko D, Yoo G, Yun S-T, Jun S-C, Chung H. Bacterial and fungal community composition across the soil depth profiles in a fallow field. Journal of Ecology and Environment. 2017;41(1):34.

71. Kuffner M, Hai B, Rattei T, Melodelima C, Schloter M, Zechmeister-Boltenstern S, et al. Effects of season and experimental warming on the bacterial community in a temperate mountain forest soil assessed by 16S rRNA gene pyrosequencing. FEMS microbiology ecology. 2012;82(3):551–62.

72. Aoki Y, Fujita K, Shima H, Suzuki S. Survey of annual and seasonal fungal communities in Japanese Prunus mume orchard soil by next-generation sequencing. Advances in Microbiology. 2015;5(13):817.

73. Kim CS, Nam JW, Jo JW, Kim S-Y, Han J-G, Hyun MW, et al. Studies on seasonal dynamics of soil-higher fungal communities in Mongolian oak-dominant Gwangneung forest in Korea. Journal of Microbiology. 2016;54(1):14–22.

74. Voříšková J, Brabcová V, Cajthaml T, Baldrian P. Seasonal dynamics of fungal communities in a temperate oak forest soil. New Phytologist. 2014;201(1):269–78.

